# Continuum approximations for lattice-free multi-species models of collective cell migration

**DOI:** 10.1101/119586

**Authors:** Oleksii M Matsiaka, Catherine J Penington, Ruth E Baker, Matthew J Simpson

## Abstract

Abstract

Cell migration within tissues involves the interaction of many cells from distinct subpopulations. In this work, we present a discrete model of collective cell migration where the motion of individual cells is driven by random forces, short range repulsion forces to mimic crowding, and longer range attraction forces to mimic adhesion. This discrete model can be used to simulate a population of cells that is composed of *K* ≥ 1 distinct subpopulations. To analyse the discrete model we formulate a hierarchy of moment equations that describe the spatial evolution of the density of agents, pairs of agents, triplets of agents, and so forth. To solve the hierarchy of moment equations we introduce two forms of closure: (i) the mean field approximation, which effectively assumes that the distributions of individual agents are independent; and (ii) a moment dynamics description that is based on the Kirkwood superposition approximation. The moment dynamics description provides an approximate way of incorporating spatial patterns, such as agent clustering, into the continuum description. Comparing the performance of the two continuum descriptions confirms that both perform well when adhesive forces are sufficiently weak. In contrast, the moment dynamics description outperforms the mean field model when adhesive forces are sufficiently large. This is a first attempt to provide an accurate continuum description of a lattice-free, multi-species model of collective cell migration.

## 1. Introduction

*In vivo* cell migration involves many different cell types interacting with each other. For example, tumour invasion involves malignant cancer cells moving through normal surrounding tissues (Weinberg, 2009). Interactions between different cell types are also captured in certain *in vitro* experiments, such as the migration of malignant melanoma cells, which is thought to be enhanced when these cells are moving amongst skin cells (Eves et al., 2003). Multiple species of cells can also be created in experiments where some subpopulation of cells, amongst an otherwise identical subpopulation, are labelled and tracked over time (Simpson et al. 2006; Simpson et al., 2007). While some mathematical models explicitly account for interactions between different subpopulations of cells (Painter and Sherratt, 2003), most mathematical models deal with a single population of cells only (Sherratt and Murray, 1990; Maini et al., 2004).

A common approach to modelling cell migration is to use a lattice-based random walk model. This approach captures details of the motion of individual cells, which is attractive because this kind of information can be linked to time lapse images from experiments. The continuum-limit description of such a lattice-based model can also be used to study the group behaviour. Although some previous lattice-based models account for interactions between different types of cells (Simpson et al., 2009; Penington et al., 2011), these lattice-based models are unrealistic because real cells do not move on regular lattice-based structures. Other limitations of lattice-based models include restrictions on cell size. For example, the diameter of a typical melanoma cell is approximately 18 *μm* (Treloar et al., 2013) whereas the diameter of a typical skin cell is approximately 25 *μm* (Simpson et al., 2013). In a model with both types of cells present, it is not possible to accommodate these differences in cell size if we use a standard lattice-based approach where each cell occupies a single lattice site (Binder and Simpson, 2016).

To address these limitations, we define a lattice-free model that can be used to describe the migration of a population of cells that is composed of many potentially distinct subpopulations. We adopt a modelling framework that is an extension of previous approaches by Newman and Grima (2004) and Middleton and co-workers (2014). The work by Newman and Grima considered a stochastic model of individual cell migration, with chemotactic effects, and they described the continuum limit using a Langevin formulation. The work of Newman and Grima (2004) was then extended by Middleton and co-workers (2014) who also considered a stochastic model of individual cell migration in terms of a Langevin formulation, however they considered both a traditional mean field continuum approximation as well as a more sophisticated moment closure continuum approximation that accounts for the spatial and temporal dynamics of pairs of agents. A key feature of both these previous models is that they are appropriate for studying the collective migration of a single populations of cells. However, many practical problems in development and disease progression involves multiple interacting subpopulations of cells. Therefore, the main aim of the current study is to develop a discrete model of collective migration where the total population of cells consists of an arbitrary number of interacting subpopulations. Our discrete model incorporates random cell motility, adhesion between cells and finite size effects (crowding). We allow for differences in cell size, cell motility and cell adhesion between the different subpopulations. In addition to producing stochastic realisations of the discrete model, we also analyse the continuum limit using both a standard mean field approximation and a more sophisticated moment dynamics approximation. Comparing averaged behaviour from the discrete simulations with the solution of the continuum models confirms that the mean field approach can be inaccurate when adhesion is sufficiently strong. This is important because almost all mathematical models of collective cell migration invoke the mean field approximation (Sherratt and Murray, 1990; Painter and Sherratt, 2003; Maini et al., 2004).

This manuscript is organised in the following way. In Section 2 we describe the discrete model. In Section 3.1, we analyse the discrete model, showing how we can obtain a continuum description of the average behaviour of the discrete model. In particular, we focus on two different continuum descriptions: (i) a mean field approximation; and (ii) a higher-order moment dynamics approximation. Results in Sections 3.2-3.3 compare solutions of both continuum approximations and averaged discrete results for problems involving one and two interacting subpopulations, with additional comparisons presented in the Supplementary Material. In Section 3.4 we investigate how the accuracy of the MFA and KSA approximations depends on the choice of model parameters. Finally, in Section 4, we summarise our work and highlight opportunities for future investigation.

## 2. Discrete model

We consider a population of *N* cells that is composed of an arbitrary number of subpopulations, *K* ≥ 1. Illustrative schematics showing interactions between individuals in a population with *K* = 1 and *K* = 2 subpopulations are given in Figure 1(a)-(b), respectively.

**Fig. 1.**
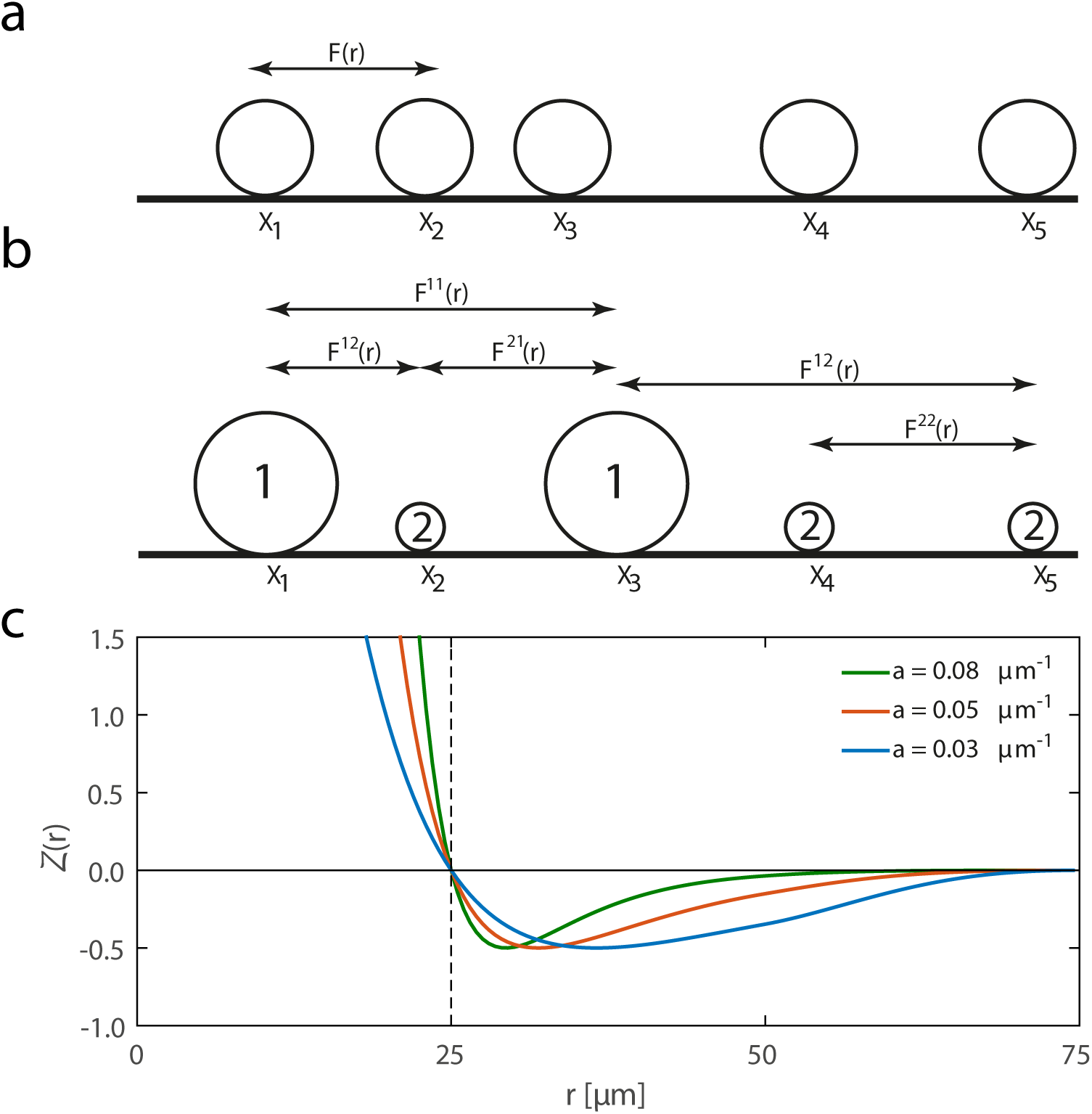
(a)-(b) Representative plot of single- and multi-species systems of cells, respectively. In (a) we show the intraspecies force, *F*(r), and in (b) we show both intraspecies forces, *F* **^11^**(*r*) and *F***^22^**(*r*), and interspecies forces, *F* **^12^**(*r*) and *F***^21^**(*r*). Here, r is the distance between cells. (c) Dimensionless force law function *Z*(*r*), given by Equation (2.5), for various values of *a*. Here, *δ* = 25 *μm* corresponds to a typical cell diameter.

We begin by assuming that each individual cell is a point mass and that its movement can be described by an equation of motion. For simplicity, from this point on, we restrict our attention to a one-dimensional geometry, and in Section 4 we discuss how the framework can be adapted to higher dimensions. To begin describing the collective motion, we assume that the motion of each cell is governed by Newton’s second law,

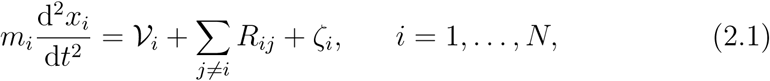
where *x_i_* is the position of the *i*_th_ cell, *m_i_* is its mass, and *R_ij_* is an interaction force between the ith and *j*_th_ cells. 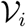 is the viscous force between the cell and the surrounding medium, and *ζ _i_* is the stochastic force associated with random Brownian motion. According to Stokes’ law, the viscous force on a small spherical particle moving in a viscous fluid is given by

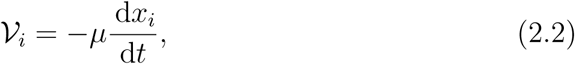
where *μ* > 0 is the drag coefficient. If we neglect inertial forces and invoke Stokes’ law (Middleton et al., 2014), we arrive at a system of Langevin stochas tic differential equations (SDEs) given by

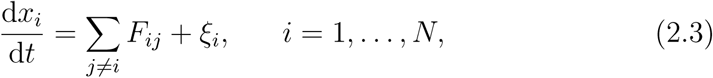
where *R_ij_*. = *μ F_ij_*. and ζ*_i_*. = *μ*ξ.

In summary, according to Equation (2.3), the collective migration of cells is determined by a balance between cell-to-cell interactions (short-range crowding and longer range adhesion), stochastic forces, and viscous forces. Collective cell migration that is driven by unbiased stochastic forces is thought to be relevant in many applications, such as collective cell spreading in many single-species in *vitro* experiments (Simpson et al., 2013). Therefore, we focus on unbiased stochastic forces by sampling *ξ_i_* from a Gaussian distribution with zero mean and zero auto-correlation (Middleton et al. 2014).

It is biologically reasonable to model the interaction forces between cells, *F_ij_*, to have different amplitudes for subpopulations of cells. This is relevant if we wish to specify different adhesion forces between different subpopulations (Steinberg, 1996). For simplicity, we assume *F_ij_* = *F_ji_*, and we specify the interaction force to be

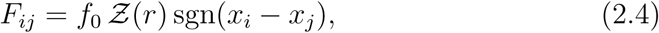
where *f***_0_** is the dimensional amplitude of the interaction force, 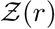 is the dimensionless force law function that depends on the separation distance, and *r* = |*x_i_* − *x_j_*|. The function sgn is the *signum* function. The particular choice of 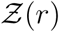 depends on phenomenological cellular behaviour we wish to model. Several force laws have been suggested, including a linear spring model (Murray et al., 2009) and non-linear force laws such as Morse (Middleton et al., 2014) or Lennard-Jones (Jeon et al., 2010) potentials. In this work we adopt a modified Morse potential force law, so that

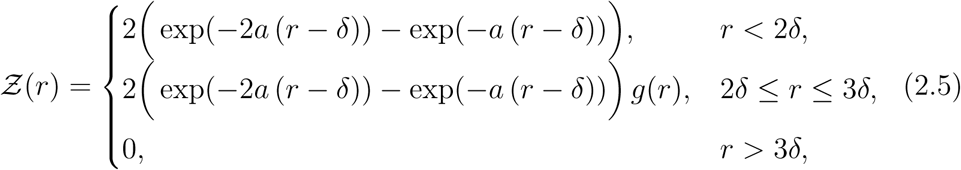
where *a* is a parameter that controls the shape of the force function, as illustrated in Figure 1(c), and *δ* is the cell diameter. The distance *r* = *δ* corresponds to the case where two cells are just in contact with each other. In this model we have 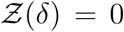 and 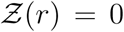 for *r* > 3 *δ*. Introducing the cell diameter *δ* in Equation (2.5) allows us to more realistically model the behaviour of multi-species populations of cells with different diameters.

Equation (2.5) incorporates the Tersoff cut-off function, *g*(*r*) (Tersoff, 1988), to capture a finite interaction range between cells. This function is given by

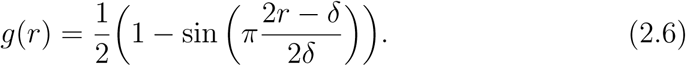

The interaction range has been chosen to be 3*δ* (Srinivas et al., 2004).

A representative plot of 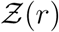 for different values of *a* is given in Figure 1(c). The force function consists of two regimes: short-range repulsion and longer range attraction. The repulsive term mimics crowding effects while the attractive tail models adhesion. While all of the results presented in this work are for this particular choice of force law, it is straightforward to incorporate other choices of 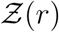.

## 3. Results and discussion

### 3.1. Mathematical model for an arbitrary number of subpopulations

We consider a total population of *N* cells that come from *K* subpopulations of cells, so that 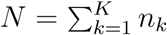, where *n_k_* is the number of cells in subpopulation *k*. This framework can be used to model both situations where each subpopulation is distinct (Eves et al. 2003) and situations where each subpopulation is composed of tagged, but otherwise identical cells (Simpson et al. 2006; Simpson et al. 2007). In addition, these distinct subpopulations may differ in many ways, such as differences in diameter, motility rates, or interaction forces and they can be arbitrarily arranged in space.

We define the one-cell probability density function (PDF), 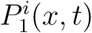, as the probability that the position of cell *i* is in the small neighbourhood [*x, x* + *dx*] at time *t*. Similarly, we define the two-cell PDF, 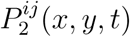, as the probability that cells *i* and *j* lie in [*x*,*x* + *dx*] and [*y*,*y* + *dy*], respectively, at time *t*. At present, we do not specify which of the subpopulations these cells belong to.

Given that the motile behaviour of cells is governed by Equation (2.3), we can relate the PDFs to the position of cells as follows (van Kampen, 1975),

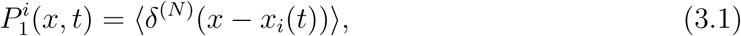

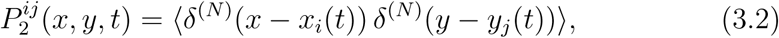
where *x*_i_(*t*) and *y*_j_(*t*) are the positions of cells given by Equation (2.3). The angled brackets indicate an average over a sufficiently large number of identically prepared initial conditions and a sufficiently large number of realisations of the stochastic force. Further background explanation about Equations (3.1) and (3.2) is given in the Supplementary Material.

The time evolution of 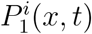 is governed by a Fokker-Planck equation (Supplementary Material),

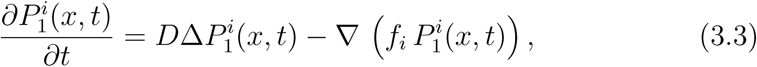
which describes the motion of particles under the influence of random forces, proportional to the diffusivity, *D*, and directed drift forces, *f_i_*. The force *f_i_* acting on cell *i* in subpopulation *l* may be expressed as the sum of two types of forces: intraspecies forces exerted by other members of subpopulation *l*, and interspecies forces exerted by cells from all other subpopulations, giving

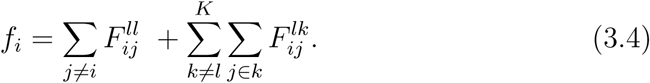

Combining Equations (3.1), (3.3) and (3.4), and taking the convolution of the

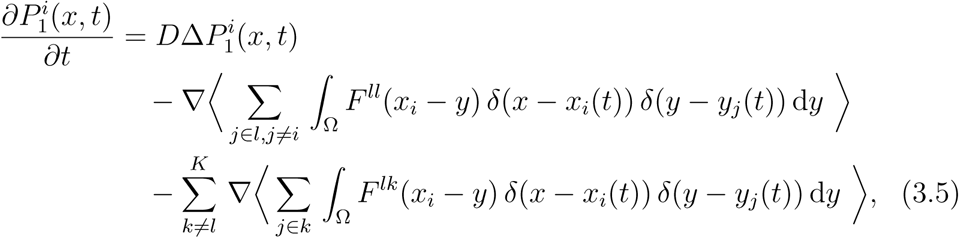
where Ω denotes the domain. The second and third terms on the right hand side of Equation (3.5) are advection terms that incorporate intraspecies and interspecies forces, respectively. Combining Equations (3.2) and (3.5), and interchanging summation and integration, we obtain

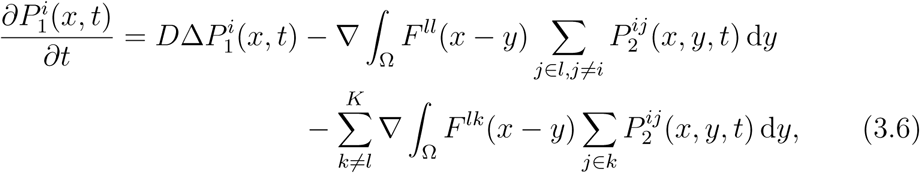
where, from this point forward, we drop the subscript *i* on *x_i_*.

To make the transition from individual level behaviour in a discrete simulation to the population level dynamics, we define the following quantities,

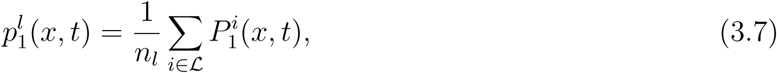

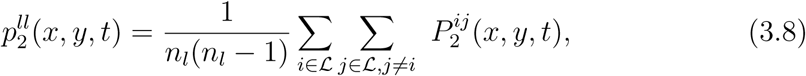

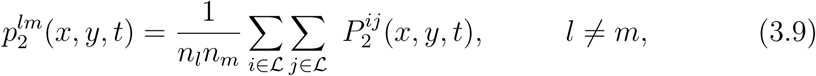
where 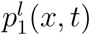 is the normalised one-cell density distribution of subpopulation *l*, 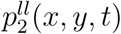 is the density-density correlation function that captures intraspecies correlations, and plm 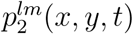 is the density-density correlation function that captures interspecies correlations.

To proceed, we sum over the index *i* in Equation (3.6) and apply the definitions given in Equations (3.7)-(3.9). We repeat this procedure *K* times for each subpopulation to yield a system of *K* non-linear integro partial differential equations (IPDEs), that can be written as

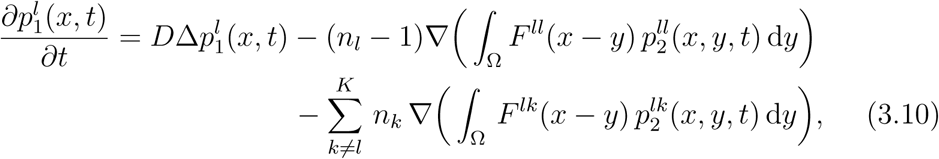
for each subpopulation *l*. We define the PDF of the total population of *N* cells as a weighted sum of the individual distributions,

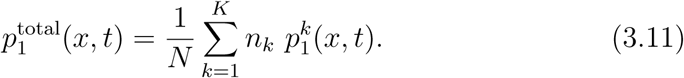

Equation (3.10) shows that the evolution of 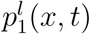 depends on 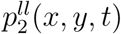. To derive an evolution equation for 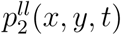we begin with the two-cell Fokker-Planck equation,

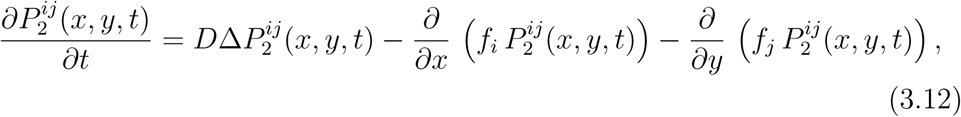
where cells *i* and *j* both belong to subpopulation *l*. The forces *f_i_* and *f_j_*, applied to cells *i* and *j*, can be written as the sum of intraspecies and interspecies forces. For example, the force on an arbitrary cell *z* can be written as

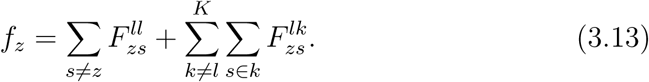

Adopting the interaction force law, Equation (2.4), using the definition of the two-cell PDF as given by Equation (3.2), and evaluating the required convolutions, we can rewrite Equation (3.12) as

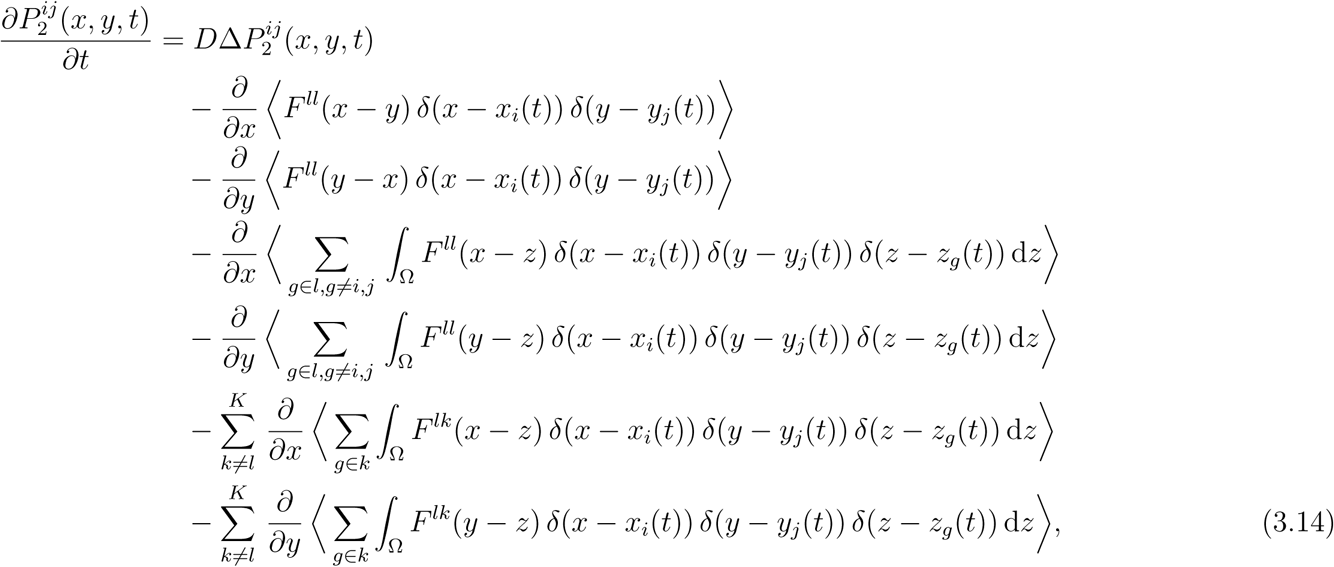
where the second and third terms on the right hand side of Equation (3.14) represent interactions between cells *i* and *j*, the fourth and fifth terms on the right hand side of Equation (3.14) represent interactions between cells i and j and other cells within subpopulation *l*, and the sixth and seventh terms on the right hand side of Equation (3.14) represent interactions between cells *i* and *j* and cells in other subpopulations.

The three-particle normalised density functions can be defined as,

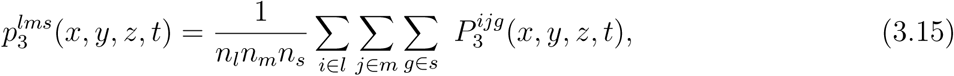

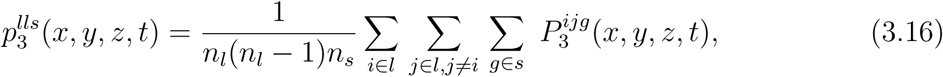

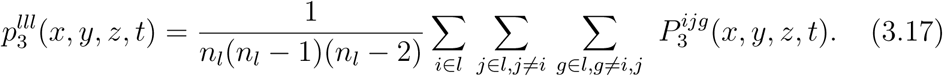

We therefore require a definition for the three particle PDF, 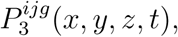, similar to Equation (3.2),

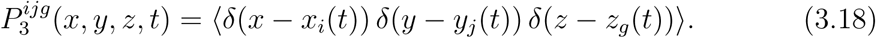

To proceed we divide Equation (3.14) by *n_l_*(*n_l_*-1), and combine Equations (3.14)-(3.18), summing over the indices i and j, to obtain an expression for the evolution of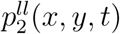,

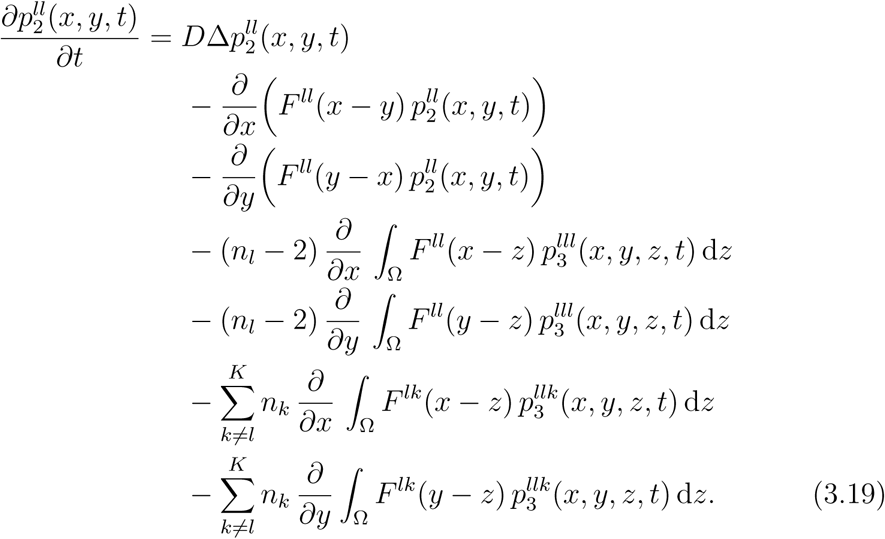

The total system of equations governing the evolution of the density-density correlation functions for *K* subpopulations consists of *K* equations in the form of Equation (3.19), and *K*! equations for the interspecies density-density correlation functions,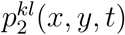.

This procedure for deriving evolution equations for the density and density-density correlation functions can be repeated to yield a hierarchy of *N* − 1 systems of IPDEs and a system of Fokker-Planck equations that govern the *N*-level density. At each level *d*, where *d* ϵ [1, *N*], the d-density function *p_d_* depends on the next order, *p_d_*_+_*_1_*. This means that the full hierarchy of equations is, in general, both analytically and numerically intractable. Therefore, we must invoke some approximations to proceed, and we will now discuss two different approximations.

#### 3.1.1 Mean field approximation

The simplest way to approximate the hierarchy is to truncate it at the first level by writing the density-density correlation function in terms of the one-cell density functions (Baker and Simpson, 2010),

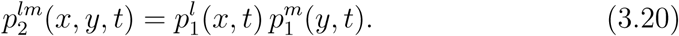

This approximation, often called the Mean Field Approximation (MFA), implies that the probability of finding one cell at [*x, x* + *dx*] at time *t* is independent of the probability of finding another cell at [*y*,*y* + *dy*] at the same time. MFA-based equations are, by far, the most popular way to describe collective cell migration (Sherratt and Murray, 1990; Painter and Sherratt, 2003; Maini et al., 2004).

We now present MFA equations for the cases relevant to both monoculture (*K* = 1) and co-culture (*K* = 2) experiments. First, for *K* =1, substituting Equation (3.20) into Equation (3.10), we obtain

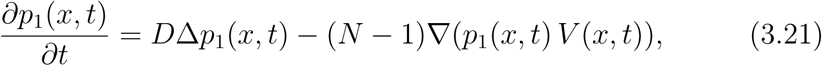
Where

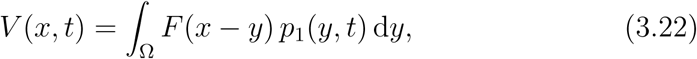
is the velocity field induced by interactions between cells. Second, for *K* = 2, the MFA leads to two coupled equations,

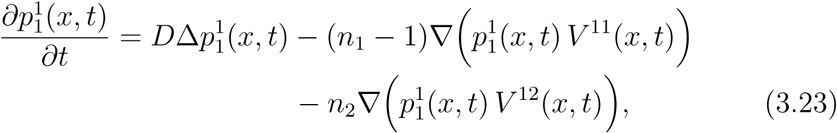

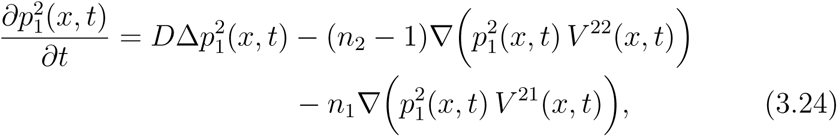

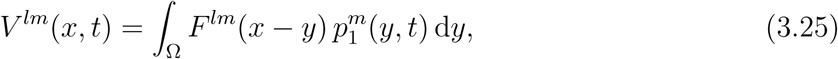
where indices *l, m* = 1, 2.

#### 3.1.2. Moment dynamics approximation

A more sophisticated approach is to use a closure relation to write for the three particle correlation function in terms of the two-particle correlation function (Baker and Simpson, 2010; Middleton et al., 2014). A commonly-used closure relations is the Kirkwood superposition approximation (KSA) (Kirkwood, 1935), which can be written as

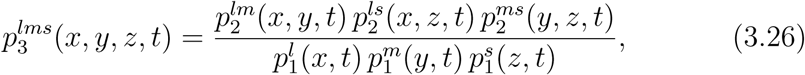
where the subpopulations *l, m* and *s* are not necessarily distinct.

For monoculture experiments with *K* = 1, the KSA continuum model can be written as

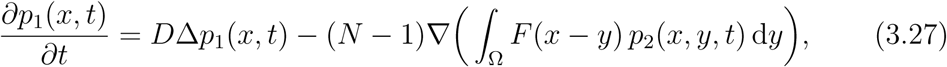

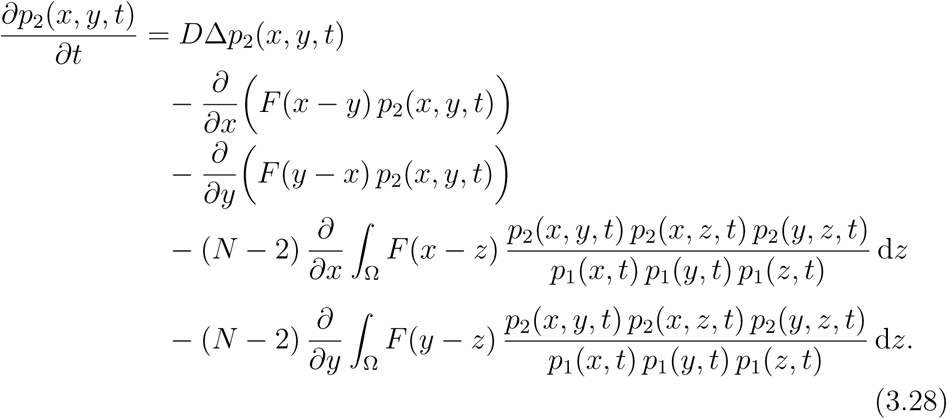

It is useful to note that there is more than one way to solve a problem with *K* =1 using the KSA framework. One approach would be to solve Equations (3.27) and (3.28) simultaneously. However, it is more computationally efficient to solve Equation (3.28) to give *p*_2_(*x*,*y*,*t*), and then to obtain *p*_1_(*x*,*t*) by numerical integration

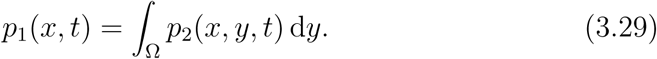

For co-culture experiments with *K* = 2, the KSA continuum model can be written as

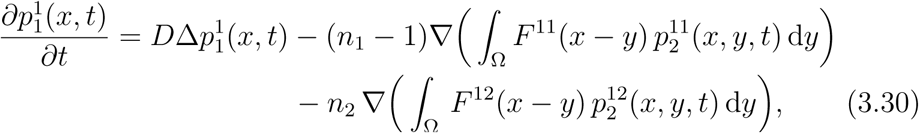

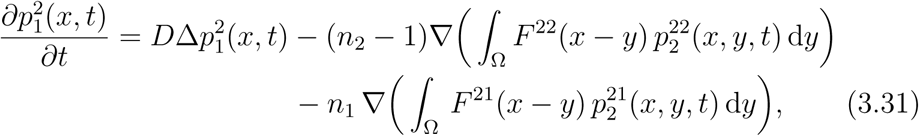

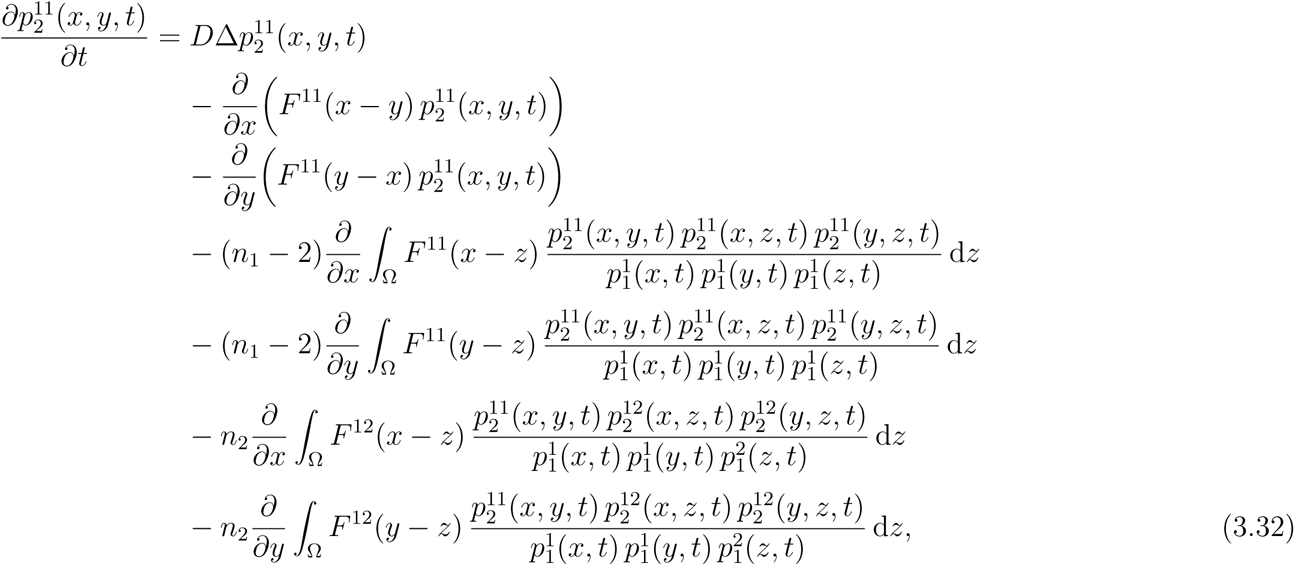

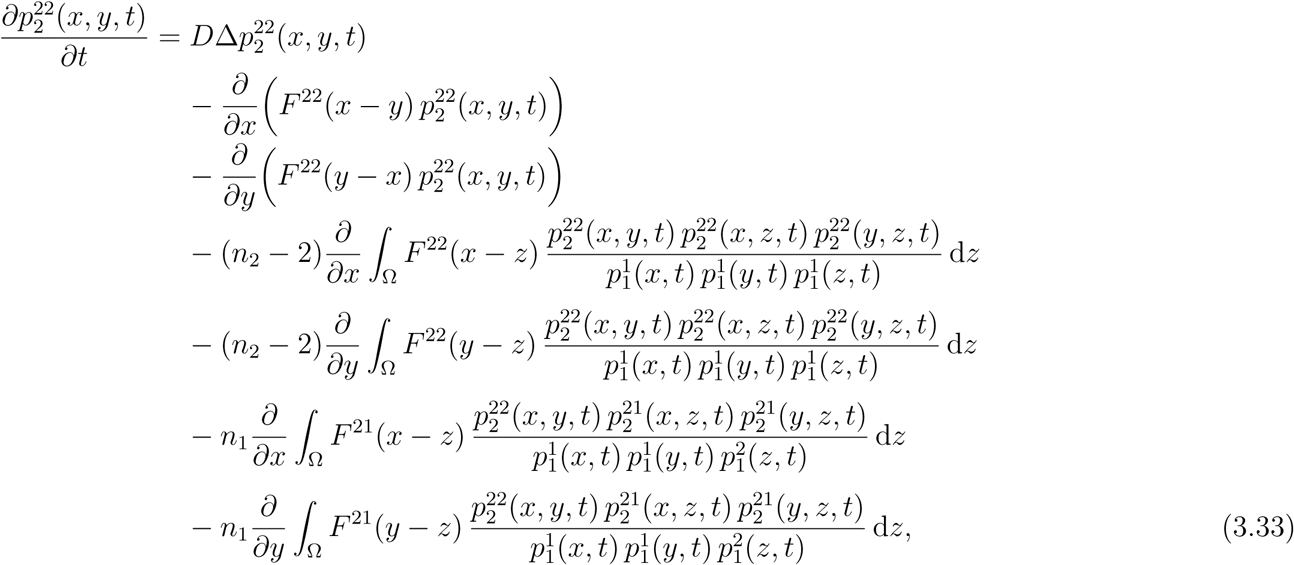

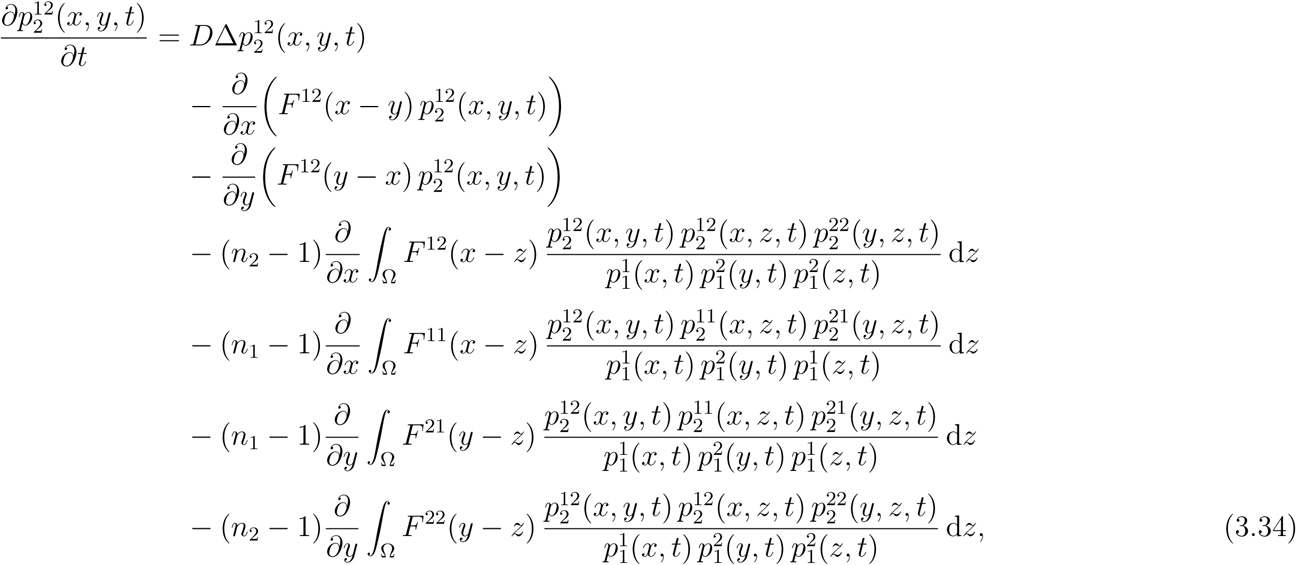

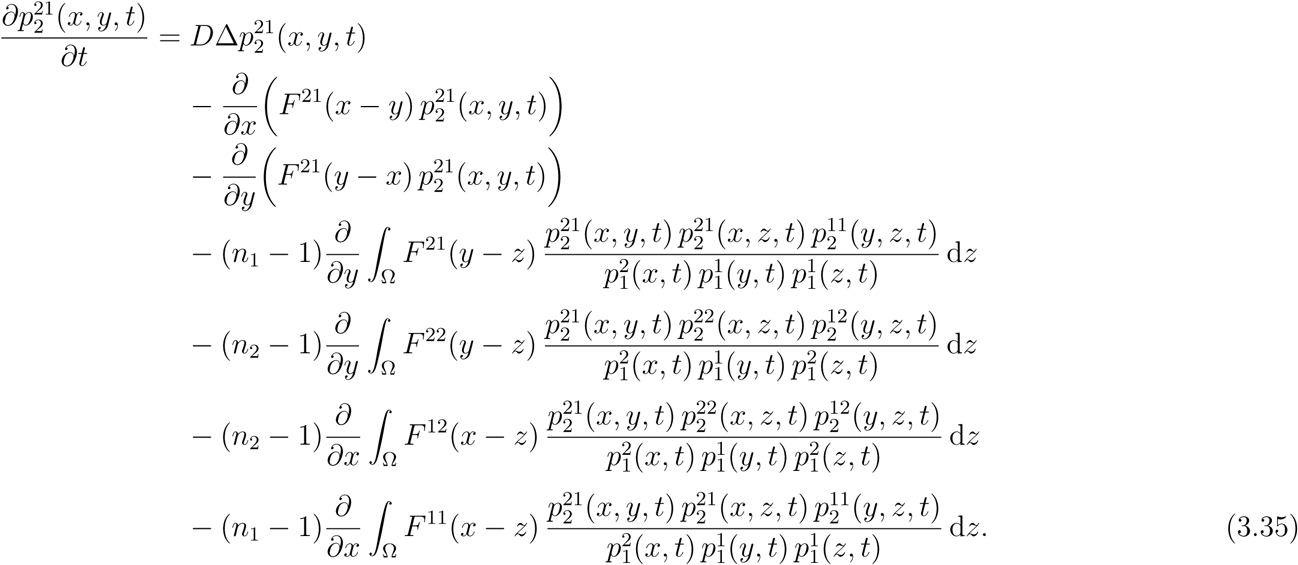

Again, there are multiple strategies for solving the KSA equations when *K* = 2. Here, we solve Equations (3.32) and (3.33) to give 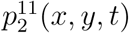 and 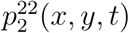, respectively. Using these results we calculate 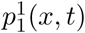 and 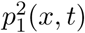 by numerical integration, similar to Equation (3.29). To obtain 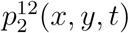 and 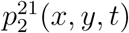, we use 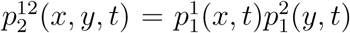 and 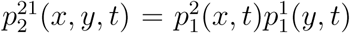, respectively.

Now that we have documented both the MFA and KSA continuum approximations for both single species monoculture (*K* = 1) and two-species co-culture (*K* = 2) experiments, we will now solve these governing equations for both cases and compare results with averaged data from discrete simulations.

### 3.2 Application to monoculture experiments, K =1

We first consider the situation where we have one population of cells, *K* = 1. In all of our numerical results we always fix the diffusivity to be *D* = 300 *μ*m^2^h^-1^ (Treloar et al. 2013). To emphasize the importance of non mean-field effects, all simulation results in the main paper involve strong adhesion, where *f***_0_** is sufficiently large. This situation is relevant when we apply our model to mimic the collective migration of epithelial cells (Treloar et al., 2013). In contrast, if the models are applied to deal with the collective migration of mesenchymal cells, without significant adhesion (Simpson et al., 2013), then additional results in the Supplementary Material document with reduced *f***_0_** are more relevant.

Since we consider unbiased random forces, we sample *ξ_i_* from a Gaussian distribution with zero mean and zero auto-correlation

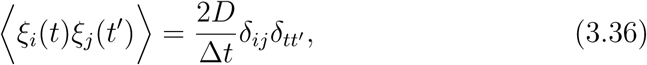
which is a white noise limit (Supplementary Material). The variance of *ξ _i_* is given by

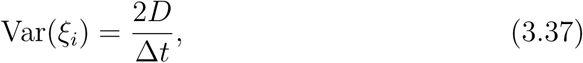
where ∆*t* is the duration of the time step used in the discrete simulations.

The initial distribution of cells in the monoculture simulations is given by

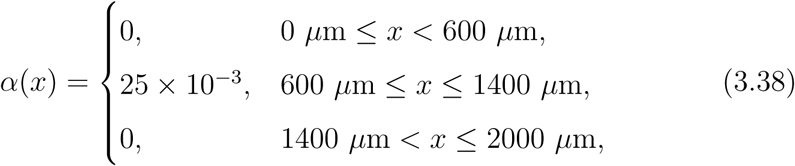
on 0 ≤ *x* ≤ 2000 *μ*m, which is a typical length scale for an *in vitro* cell migration experiment (Jin et al. 2016). Here, α(*x*) is a function of position, and we sample from this function to define the initial distribution of cells in the discrete simulations. This initial distribution corresponds to a confined group of cells in the centre of the domain. When presenting results from simulations we refer to both the dimensional density of cells, *p*_1_(*x*,*t*) [cells/*μ*m], as well as the non-dimensional density of cells relative to the carrying capacity density, *p*_1_(*x,t*)/*C*, where *C* is the carrying capacity density that is given by *C* = *Νδ* /*L*, where *N* is the maximum number of cells of diameter *δ* that can be distributed along a domain of length *L* without compression. Periodic boundary conditions are imposed for all simulations.

To solve the MFA model, we set *p*_1_(*x*, 0) = *α*(*x*), and to solve the KSA model, we note that since cells are randomly placed according to Equation (3.38), there are no spatial correlations in the initial positions of the cells. Therefore, the initial conditions for the KSA model are given by *p*_1_(*x*, 0) = *α*(*x*) and *p*_2_(*x*,*y*,0) = *α*(*x*) *α*(*y*). With this information, Equations (3.21) and (3.28) are solved using the method of lines with spatial and temporal discretisations chosen to be sufficiently fine that the numerical solutions are grid independent. The discrete model, Equation (2.3), is numerically integrated using a fourth order Runge-Kutta (RK4) method (Press et al., 2007) and density distributions are obtained by considering a large number of identically prepared simulations. Results in Figure 2 compare numerical solutions of the MFA and KSA continuum descriptions with averaged data from discrete simulations. Snapshots of the discrete simulations are shown in Figure 2(a)-(b). A comparison of the ensemble averaged data and the solution of the MFA and KSA models are given in Figure 2(c) and Figure 2(e), respectively. To clearly compare the performance of the MFA and KSA models near the position of the spreading profile, we show a magnified region of the profiles in Figures 2(d) and Figure 2(f).

**Fig. 2.**
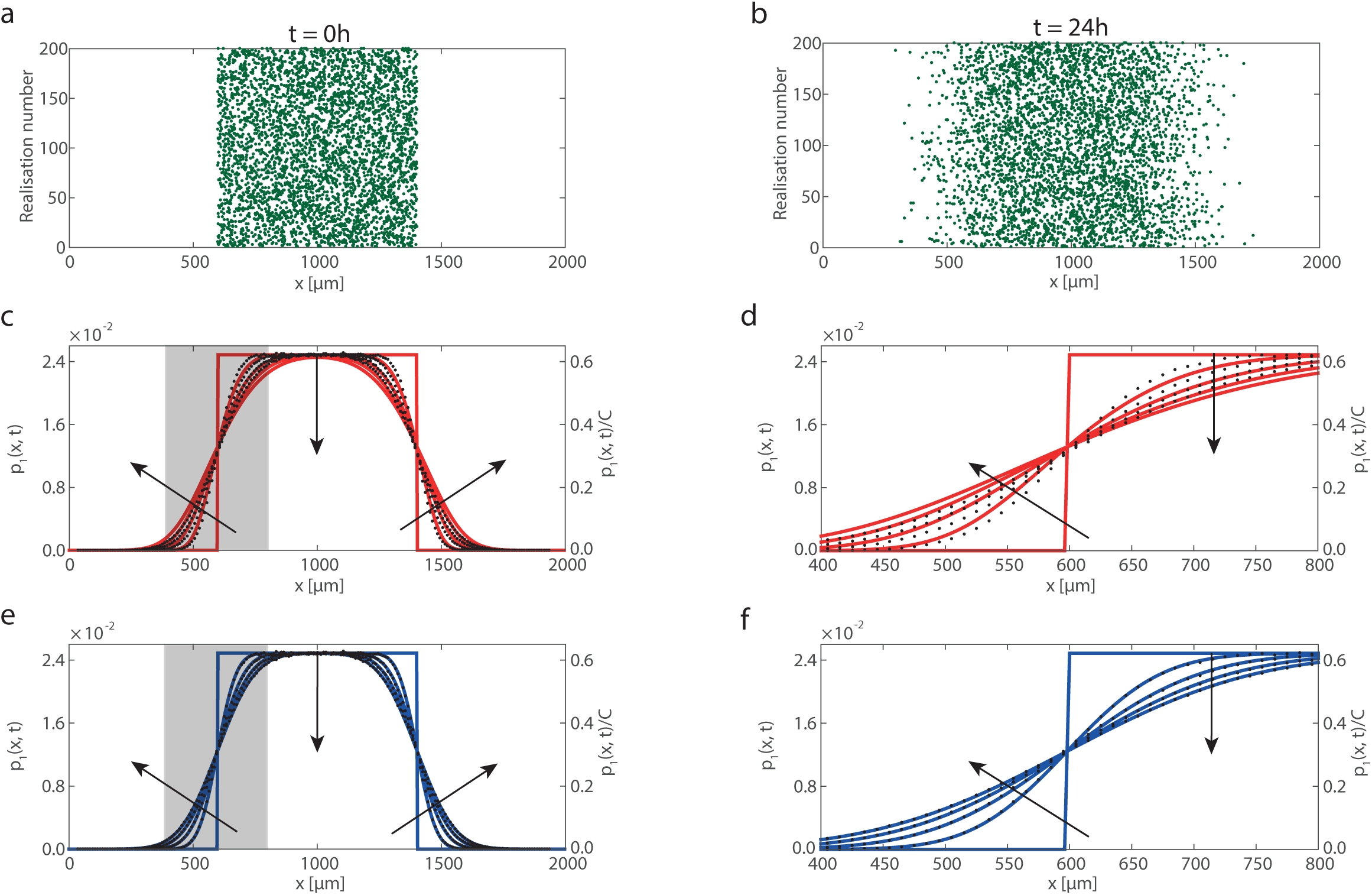
Comparison of ensemble averages of stochastic simulations and solutions of the MFA and KSA continuum models for a single-species population of cells on a one-dimensional domain with 0 ≤ *x* ≤ 2000 *μ*m. Snapshots in (a)-(b) show 200 realisations of the discrete model at *t* = 0 and *t* = 24 hours, respectively. The population of cells (green) initially occupies the central region, which is 800 *μ*m wide, and has an initial density 25 × 10^−3^ cells/*μ*m. Results in (c)-(f) show the cell density profiles obtained using an ensemble of 5 × 10^5^ simulations (black dots) with binsize of 10 *μ*m. These results are compared to solutions of the MFA model, Equation (3.21) (red lines), and the KSA model, Equation (3.28) (blue lines). Profiles are given at *t* = 0,6,12,18, and 24 h, with the arrows indicating the direction of increasing *t*. In (c)-(f) the cell density is reported in terms of the dimensional cell density, *p*_1_(*x*,*t*), as well as the dimensionless cell density, *p_1_*(*x*,*t*)*/C*, where *C* = 40 × 10^−3^ cells/*μ*m. Equation (2.3) is integrated with Δ*t* = 5 × 10^−2^ h, Equation (3.21) is integrated with Δ*x* = 4*μ*m and Δ*t* = 10^−2^ h, and Equation (3.28) is integrated with Δ*x* = Δ*y* = 4 *μ*m and Δ*t* = 10^−2^ h. The remaining parameters are *N* = 20, *a* = 0.08 *μ*m^-1^, *δ* = 25 *μ*m, *f*_0_ = 0.2 *μ*m/h.

In summary, we see that both the KSA and MFA models capture the overall spreading behaviour of the collective migration reasonably well, as shown in Figure 2(c) and Figure 2(e). However, when we examine the performance of MFA model more closely, as illustrated in Figure 2(d), we see that the solution of the MFA continuum model is not as steep as the discrete density data. In contrast, the performance of the KSA model, as shown in Figure 2(f), provides an improved match to the averaged discrete data. We now examine the relative performance of the MFA and KSA approaches for two multi-species problems.

### 3.3 Application to co-culture experiments, K = 2

We now consider the evolution of two types of two-species problems. These two problems involve different experimental designs. In both cases we choose the size of the cells in subpopulations 1 and 2 to be different. Here, the diameter of cells in the first subpopulation is *δ*_1_= 18 *μ*m, and the diameter of cells in the second subpopulation is *δ*_2_ = 25 *μ*m. We also introduce differing interspecies interaction parameters such as the interspecies force amplitude, 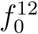, shape parameter, *a*_12_, and the interspecies diameter, *δ*_12_, which corresponds to the average radius of the different cell types.

The first experiment involves one population of cells spreading through another background population of cells, and this mimics the way that an initially confined population of tumour cells might spread through surrounding healthy tissue (Eves et al. 2003). To specify the initial condition for this problem we must describe the initial location of both subpopulations,

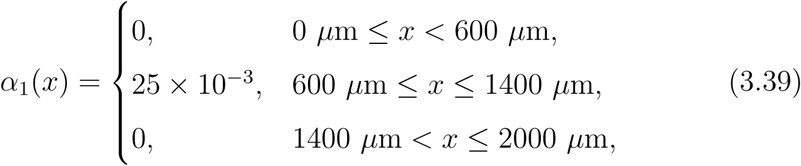

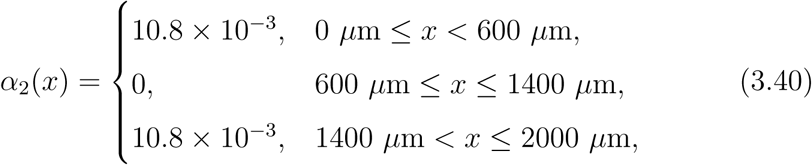
where *α*_1_(*x*) is a function of position that describes the initial location of cells from the first subpopulation, and *α*_2_(*x*) is a function of position that describes the initial location of cells from the second subpopulation. This initial condition corresponds to the situation where the region 600 ≤ *x* ≤ 1400 *μ*m is relatively densely occupied by subpopulation 1, and the remaining space is less densely populated by subpopulation 2. To initialise the discrete simulations we sample from *α*_1_(*x*) and *α*_2_(*x*), and snapshots showing 200 realisations of discrete model are given in Figure 3(a)-(c) at *t* = 0, 12 and 24 hours, showing how the two subpopulations mix.

**Fig. 3.**
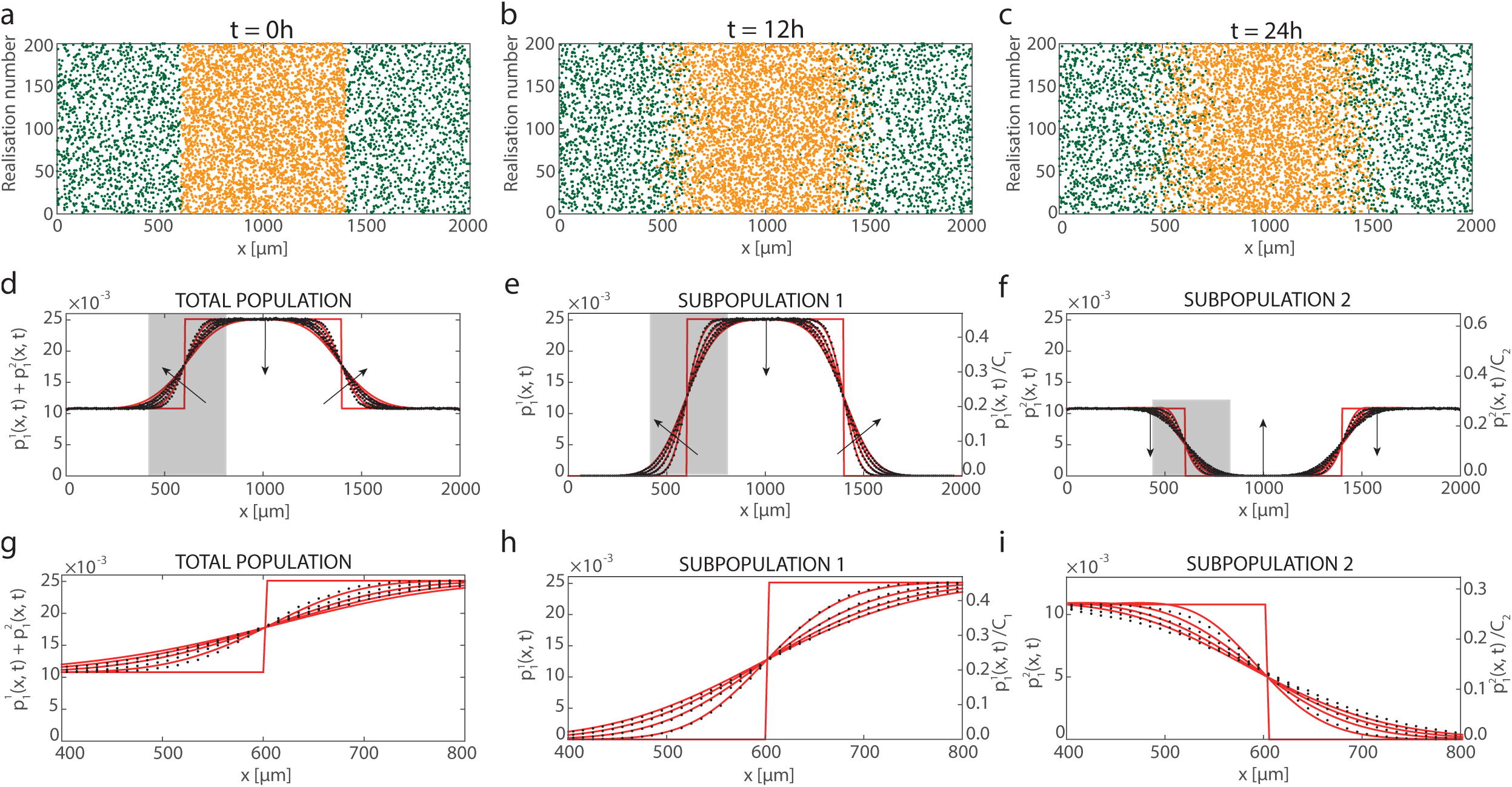
Comparison of ensemble averages of stochastic simulations and solutions of the MFA continuum model, given by Equations (3.23)-(3.24), for a two-species population of cells on a one-dimensional domain with 0 ≤ *x* ≤ 2000 *μ*m. Snapshots in (a)-(c) show 200 realisations of the discrete model at *t* = 0,12, and 24 hours, respectively. Subpopulation 1 (orange) initially occupies the central region at a density of 25 × 10^−3^ cells/*μ*m, and subpopulation 2 (green) initially occupies two outer regions at a density of 10.8 × 10^−3^ cells/*μ*m. Results in (d)-(i) show the density profiles obtained using an ensemble of 5 × 10^5^ simulations (black dots) with binsize of 10 *μ*m, and the solutions of the MFA model (red lines) at *t* = 0,6,12,18 and 24 h, with the arrows indicating the direction of increasing *t*. Results in (d)-(i) are shown in terms of the total population density, the density of subpopulation 1, and the density of subpopulation 2, as indicated. Density profiles are reported in terms of the dimensional cell densities, 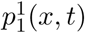 and 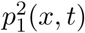, as well as the dimensionless cell densities, 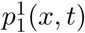/ *C*1 and 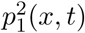/*C*_2_, where *C*_1_ = 55.5 × 10^−3^ cells/ *μ*m, *C_2_* = 40 × 10^−3^ cells/*μ*m. The MFA model is integrated with Δ*x* = 4 *μ*m and Δ*t* = 5 × 10^−3^ h. The remaining parameters are *n_1_* = 20, *n_2_* = 13, *a_1_* = 0.08 *μ*m^-1^, *a_2_* = 0.06 *μ*m^-1^, a_12_ = 0.07 *μ*m^-1^, *δ*_2_ = 18 *μ*m, *δ_2_* = 25 *μ*m, *δ*_12_ = 21.5 *μ*m, 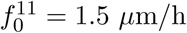, 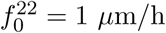, 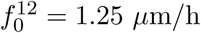.

**Fig. 4.**
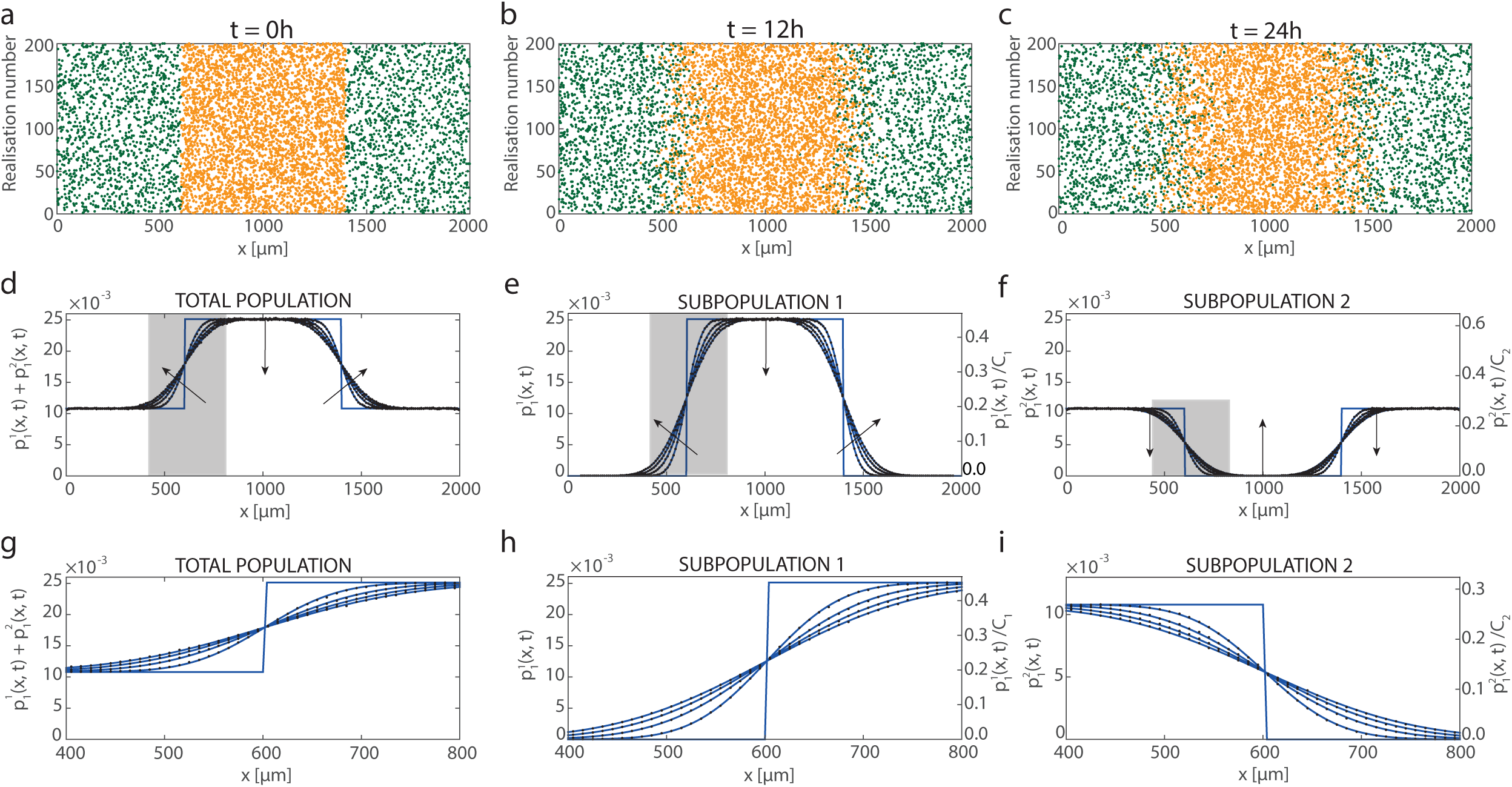
Comparison of ensemble averages of stochastic simulations and solutions of the KSA continuum model, given by Equations (3.32)-(3.33), for a two-species population of cells on a one-dimensional domain with 0 ≤ *x* ≤ 2000 *μ*m. Snapshots in (a)-(c) show 200 realisations of the discrete model at *t* = 0, 12, and 24 hours, respectively. Subpopulation 1 (orange) initially occupies the central region at a density of 25 × 10^−3^ cells/*μ*m, and subpopultation 2 (green) initially occupies two outer regions at a density of 10.8 × 10^−3^ cells/*μ*m. Results in (d)-(i) show the density profiles obtained using an ensemble of 5 × 10^5^ simulations (black dots) with binsize of 10 *μ*m, and the solutions of the KSA model (red lines) at *t* = 0,6,12,18 and 24 h, with the arrows indicating the direction of increasing t. Results in (d)-(i) are shown in terms of the total population density, the density of subpopulation 1 and the density of subpopulation 2, as indicated. Density profiles are reported in terms of the dimensional cell densities, 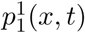 and 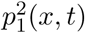, as well as the dimensionless cell densities and 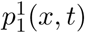/*C*_1_ and 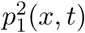/*C*_2_ where *C*_1_ = 55.5 × 10^−3^ cells/*μ*m, *C*_2_ = 40 × 10^−3^ cells/*μ*m. The KSA model is integrated with Δ*x* = Δ*y* = 4 *μ*m and Δ*t* = 5 × 10^−3^ h. The remaining parameters are *n*_1_ = 20, *n*_2_ = 13, *a*_1_ = 0.08 *μ*m^-1^, *a*_2_ = 0.06 *μ*m^-1^, *a*_12_ = 0.07 *μ*m^-1^, *δ*_1_ = 18 *μ*m, *δ*_2_ = 25 *μ*m, *δ*_12_ = 21.5 *μ*m, 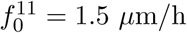,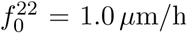,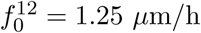

The second experiment that we consider corresponds to two initially adjacent subpopulations of cells. The initial location of both subpopulations is given

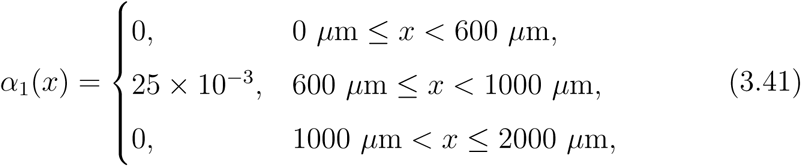

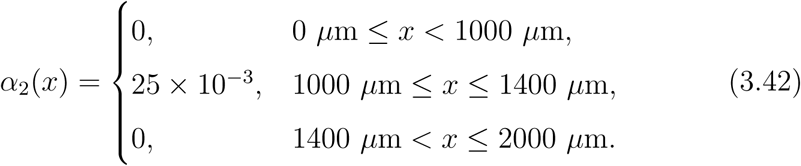

To initialise the discrete simulations we sample from *α*_1_(*x*) and *α*_2_(*x*), and snapshots showing 200 realisations of discrete model for the second initial condition are given in Figure 5(a)-(c) at *t* = 0, 12 and 24 hours. Here we see that the two subpopulations mix near *x* = 1000 *μ*m. Furthermore, we also see that both subpopulations spread into the initially vacant surrounding regions.

**Fig. 5.**
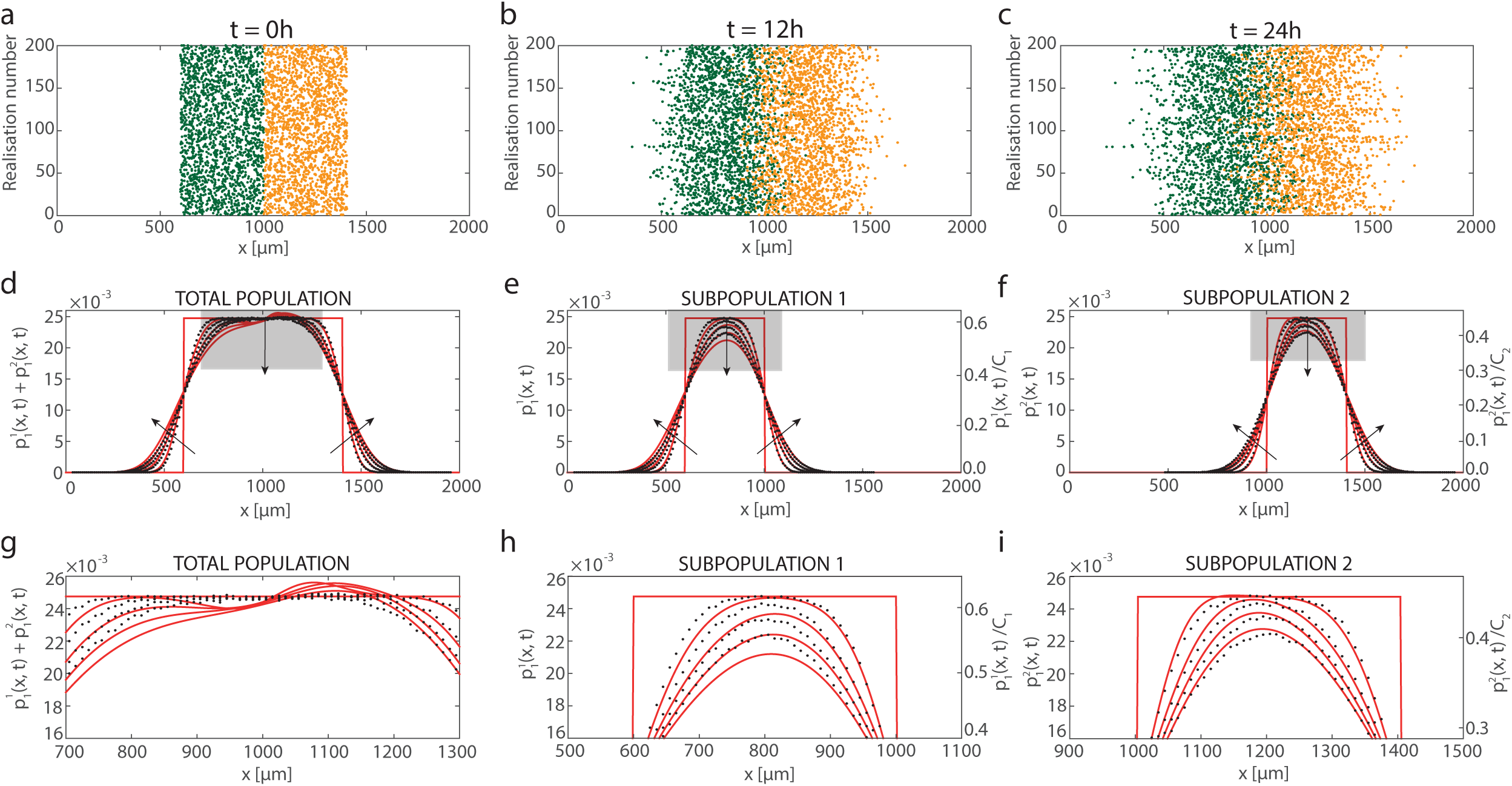
Comparison of ensemble averages of stochastic simulations and solution of the MFA continuum model, given by Equations (3.23)-(3.24), for a two-species population of cells on a one-dimensional domain with 0 ≤*x* ≤ 2000 *μ*m. Snapshots in (a)-(c) show 200 realisations of the discrete model at *t* = 0,12, and 24 hours, respectively. Subpopulations 1 (green) and 2 (orange) initially occupy adjacent regions at a density of 25 × 10^−3^ cells/*μ*m. Results in (d)-(i) show the density profiles obtained using an ensemble of 5 × 10^5^ simulations (black dots) with binsize of 10 *μ*m, and the solutions of the MFA model (red lines) at *t* = 0,6,12,18 and 24 h, with the arrows indicating the direction of increasing *t*. Results in (d)-(i) are shown in terms of the total population density, the density of subpopulation 1 and the density of subpopulation 2, as indicated. Density profiles are reported in terms of the dimensional cell densities, 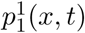 and 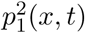, as well as the dimensionless cell densities, 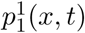/*C*_1_ and 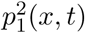/*C*_2_, where *C*_1_ = 40 × 10^−3^ cells/*μ*m, C_2_ = 55.5 × 10^−3^ cells/*μ*m. The MFA model is integrated with Δ*x* = 4 *μ*m and Δ*t* = 5 × 10^−3^ h. The remaining parameters are *n*_1_ = 10, *n*_2_ = 10, *a*_2_ = 0.06 *μ*m^-1^, *a*_2_ = 0.08 *μ*m^-1^, *a*_12_ = 0.07 *μ*m^-1^, *δ*_1_ = 25 *μ*m, *δ*_2_ = 18 *μ*m, *δ*_12_ = 21.5 *μ*m, 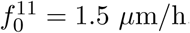, 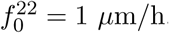, 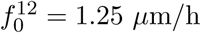

**Fig. 6.**
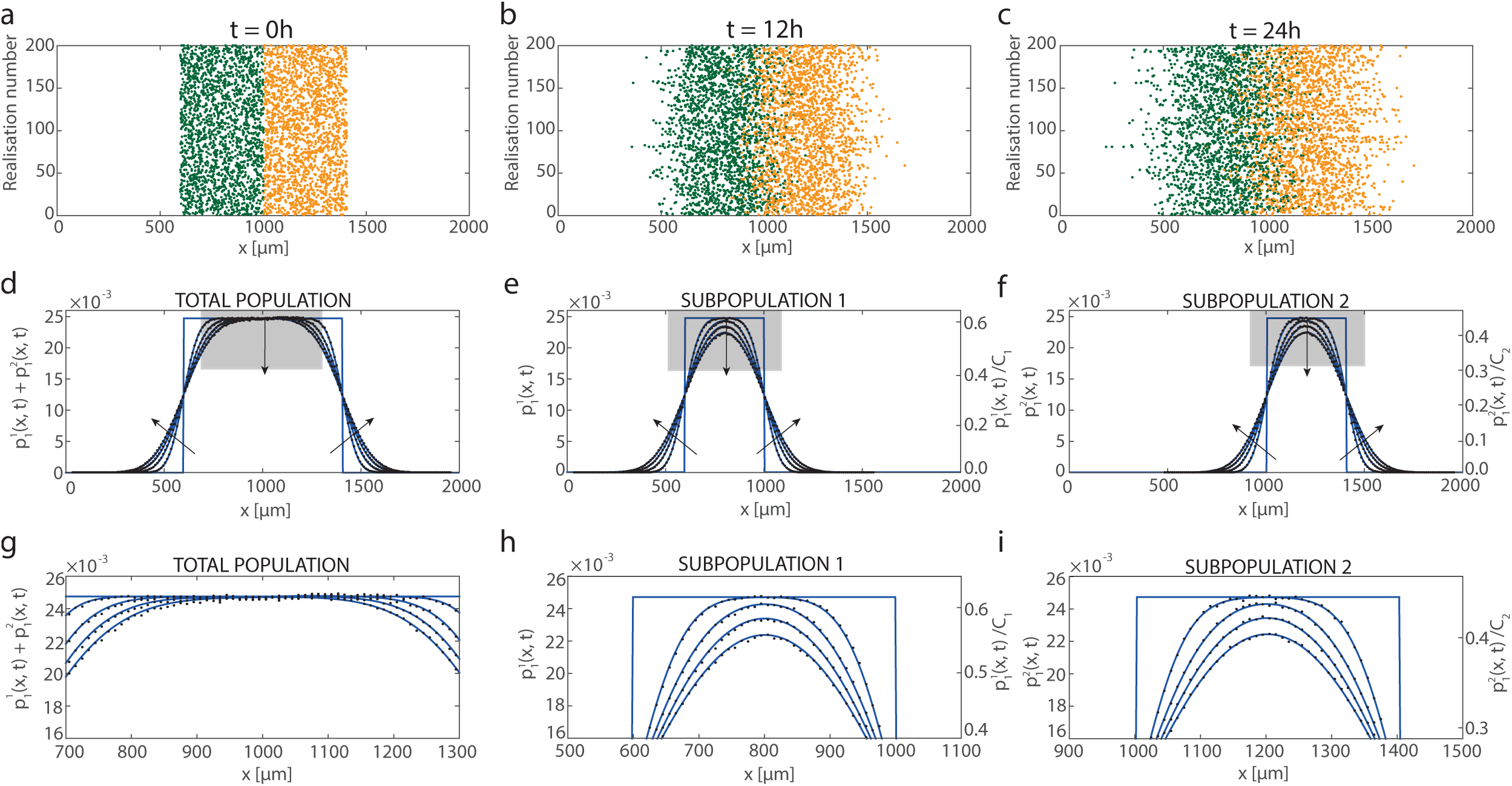
Comparison of ensemble averages of stochastic simulations and solutions of the KSA continuum model, given by Equations (3.32)-(3.33), for a two-species population of cells on a one-dimensional domain with 0 ≤ × ≤ 2000 *μ*m. Snapshots in (a)-(c) show 200 realisations of the discrete model at *t* = 0, 12, and 24 hours, respectively. Subpopulations 1 (green) and 2 (orange) initially occupy adjacent regions at a density of 25 × 10^−3^ cells/*μ*m. Results in (d)-(i) show the density profiles obtained using an ensemble of 5 × 10^5^ simulations (black dots) with binsize of 10 *μ*m, and the solutions of the KSA Equations (blue lines) at *t* = 0,6,12,18 and 24 h, with the arrows indicating the direction of increasing *t*. Results in (d)-(i) are shown in terms of the total population density, the density of subpopulation 1 and the density of subpopulation 2, as indicated. Density profiles are reported in terms of the dimensional cell densities, 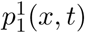and 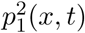, as well as the dimensionless cell densities, 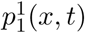 /*C*_1_ and 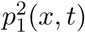/*C*_2_, where *C*_1_ = 40 × 10^−3^ cells/*μ*m, *C*_2_ = 55.5 × 10^−3^ cells/*μ*m. The KSA model is integrated with Δ*x* = Δ*y* = 4 *μ*m and Δ*t* = 5 × 10^−3^ h. The remaining parameters are n_1_ = 10, n_2_ = 10, a_1_ = 0.06 *μ*m^-1^, a_2_ = 0.08 *μ*m^-1^, a_12_ = 0.07 *μ*m^-1^, *δ*_1_ = 25 *μ*m, *δ*_2_ = 18 *μ*m, *δ*_12_ = 21.5 *μ*m, 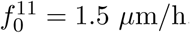,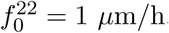,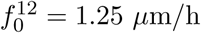

To obtain continuum results for the two-species problems, the MFA and KSA continuum models, given by Equations (3.23)-(3.24) and Equations (3.32)-(3.33), respectively, are solved using the method of lines with spatial and temporal discretisations chosen to be sufficiently fine that the numerical solutions are grid independent (Supplementary Material). Results in Figures 3 and 5 compare the performance of the MFA approach with the averaged discrete data. Since these simulations involve significant interaction forces, we see that the solution of the MFA model does not always accurately capture the details of how the subpopulations spread and interact with each other. Results in Figures 4 and 6 compare the performance of the KSA approach with the averaged discrete data. Comparing the performance of the KSA and MFA models confirms that, similar to our results for the single-species problem in Figure 2, the KSA approach outperforms the MFA model.

### 3.4 Parameter sensitivity

In this section we investigate how the accuracy of the both continuum approximations depends on the choice of the model parameters. To explore this question we re-examine the results of the first co-culture experiment, as illustrated in Figures 3-4, and we quantify how the accuracy of the KSA and MFA continuum models depends on the strength of adhesion and the ratio of the two cell sizes in the co-culture experiment. To explore this we repeat the discrete simulations and vary the force amplitude 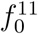, which determines strength of the cell-to-cell adhesion, as well as varying the ratio *δ*_1_/*δ*_2_. To keep our analysis as straightforward as possible, we vary these two quantities separately.

To quantify the accuracy of both the MFA and KSA continuum approximations we define the following quantities,

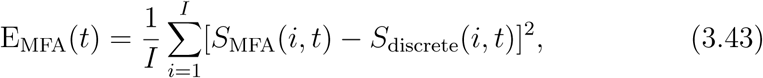

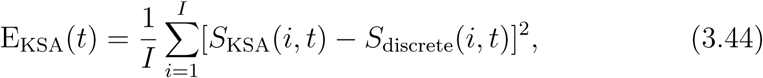
where E_MFA_(*t*) and E_KSA_(*t*) indicate mean squared error associated with the MFA and KSA approximations, respectively. The index *i* denotes the spatial node, and *I* = 200 is the total number of spatial nodes across the domain. To construct these mean squared errors we compare the total density profiles so that 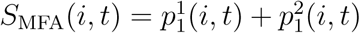is the total population density predicted by the MFA continuum approximation, 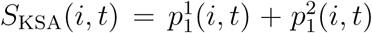 is the total population density predicted by the KSA continuum approximation, and 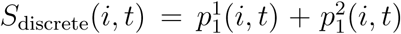 is the total population density obtained by considering an ensemble average of the discrete model.

Results in Figure 7 show E_MFA_(*t*) and E_ksA_ (*t*) as a function of *δ*_1_/*δ*_2_ and 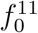. The vertical lines correspond to choices of *δ*_1_/*δ*_2_ and 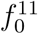 that are identical to the parameter values used to construct the results in Figures 3-4. Overall, the results in Figure 7 show three main trends: (i) for all parameter choices considered in the sensitivity analysis, the KSA approximation outperforms the MFA approximation; (ii) the accuracy of both the MFA and KSA approximations decrease with both *δ*_1_/*δ*_2_ and 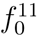; and (iii) the accuracy of both the MFA and KSA approximations is more sensitive to changes in *δ*_1_/*δ*_2_ than changes in 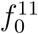 for the range of parameters considered.

**Fig. 7.**
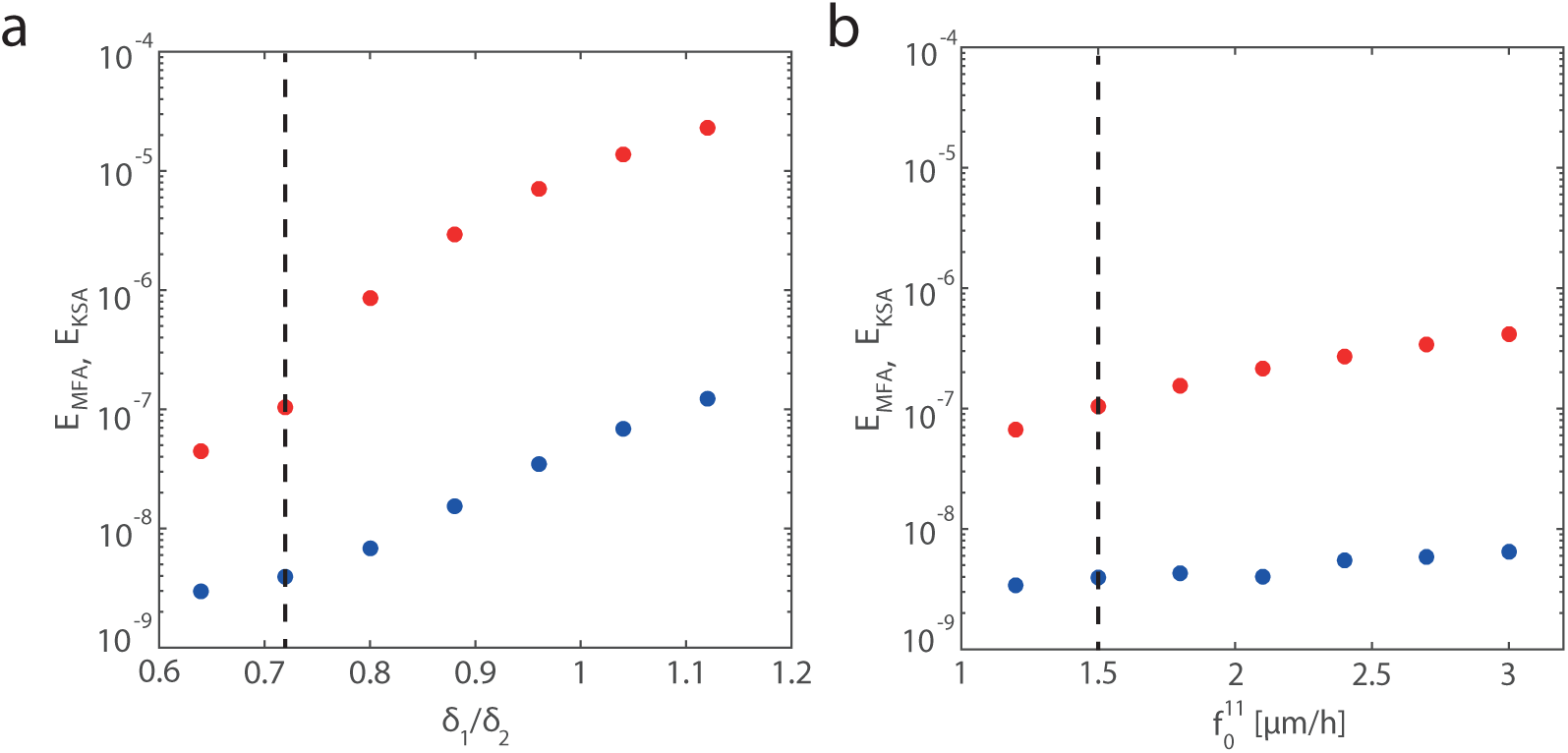
(a) Comparison of the accuracy of the MFA and KSA continuum approximations as a function of *δ*_1_/*δ*_2_ at time *t* = 24 h for the first co-culture experiment. All results in (a) correspond to *δ*_2_ = 25 *μ*m, and the ratio *δ*_1_/*δ*_2_ is varied by altering *δ*_1_. (b) Comparison of the accuracy of the MFA and KSA continuum approximations as a function of 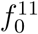at time *t* = 24 h for the first co-culture experiment. All data in (b) correspond to a fixed choice of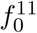. Both subfigures show E_MFA_(*t*) (red dots) and EKSA(*t*) (blue dots), and the dashed vertical line indicates the parameter values presented previously in Figures 3-4. All continuum models are solved numerically with Δ_x_ = 4 *μ*m and Δ*t* = 5 × 10^−3^ *h*.

## 4 Conclusions

In this work, we develop a discrete multi-species model of collective cell migration. Our framework is very general, and can deal with genuine multi-species problems where the subpopulations are distinct (Eves et al., 2003), as well as other types of experiments where an otherwise identical subpopulation of cells is labelled (Simpson et al., 2007). Our discrete modelling framework can include various effects such as: random unbiased stochastic motion of individual cells; short range finite size effects to account for crowding interactions; longer range adhesive forces; as well as dealing with subpopulations of cells that have different cell diameters.

To analyse the discrete model, we derive a hierarchy of continuum moment equations to describe the spatial dynamics of agents, pairs of agents, triplets of agents, and so forth. We then develop two different approximate solutions of the hierarchy of moment equations. Firstly, using the MFA, and secondly, using the KSA. We compare both continuum approximations with ensemble averages from discrete simulations.

Overall, both continuum approximations match the broad features of the discrete results reasonably well. When there is little or no adhesion, both continuum models match the averaged discrete results extremely well. However, once the adhesive force is sufficiently strong, the KSA continuum model matches the averaged discrete results much better than MFA model. This difference is the consequence of adhesion causing correlations in the positions of agents in the discrete simulations (Baker and Simpson, 2010). These effects are neglected in the MFA model, however the KSA model explicitly includes the effects of pairwise correlations, *p_2_*(*x*,*y*,*t*).

There are many potential extensions which we leave for future analysis. All our analysis has been in one-dimension, but many biological experiments are in two or three dimensions (Treloar et al., 2013; Eves et al., 2003). It is relatively straightforward to apply our continuum models to higher dimensional problems, however we choose to take the most fundamental approach here and focus on one-dimension only. As it stands, isolated individual cells in our discrete model move due to unbiased random motion. However, in many applications cells move with a bias, such as in chemotaxis (Keller and Segel, 1971). To extend our model to deal with chemotaxis we would need to introduce an evolution equation for some kind of nutrient, and to allow individual cells to move with some bias in response to the spatial gradient of the nutrient (Keller and Segel, 1971). We also note that all non-MFA results are obtained by approximately closing the system of continuum equations using the KSA, however other kinds of closure relations could also be used (Murrell et al., 2004; Frasca and Sharkey, 2016).

## Acknowledgements

This work is supported by the Australian Research Council (FT130100148, DP170100474). Computational resources are provided by the High Performance Computing and Research Support Group at QUT. We appreciate the helpful comments and suggestions from the referee.

